# A theophylline-responsive riboswitch regulates expression of nuclear-encoded genes in Arabidopsis

**DOI:** 10.1101/818633

**Authors:** Nana Shanidze, Felina Lenkeit, Jörg S. Hartig, Dietmar Funck

## Abstract

Ligand-responsive synthetic riboswitches are versatile and innovative tools for external gene regulation in pro- and eukaryotes. Riboswitches are small *cis*-regulatory RNA elements that regulate gene expression by conformational changes in response to ligand binding. In plants, synthetic riboswitches were used to regulate gene expression in plastids, but the application of synthetic riboswitches for the regulation of nuclear-encoded genes *in planta* has not been reported so far. Here we characterize the properties of a theophylline-responsive synthetic aptazyme for control of nuclear-encoded transgenes in Arabidopsis (*Arabidopsis thaliana*). Activation of the aptazyme, inserted in the 3-UTR of the target gene, resulted in rapid self-cleavage and subsequent decay of the mRNA. This riboswitch allowed reversible, theophylline-dependent downregulation of the *Green Fluorescent Protein* (*GFP*) reporter gene in a dose- and time-dependent manner. Insertion of the riboswitch into the *One Helix Protein 1* (*OHP1*) gene allowed complementation of *ohp1* mutants and induction of the mutant phenotype by theophylline. *GFP* or *OHP1* transcript levels were downregulated by maximally 90%, and GFP protein levels by 95%. These results establish artificial riboswitches as tools for externally controlled gene expression in synthetic biology in plants or functional crop design.

**One sentence summary:** Artificial, ligand-responsive RNA aptazymes are an efficient tool for dose- and time-dependent external control of nuclear gene expression in plants.

## INTRODUCTION

The discovery of riboswitches has opened the possibility to design novel RNA-based systems for external control of gene expression. Riboswitches are widely distributed in prokaryotes, where they regulate transcription or translation in response to binding of a small molecule such as a metabolite or signaling compound (Mellin and Cossart, 2015; Sherwood and Henkin, 2016). Naturally occurring riboswitches are *cis*-regulatory RNA elements that are typically formed from two domains: a ligand-binding domain (aptamer) and an output domain (expression platform) that controls gene expression through a variety of mechanisms. Riboswitches are often located downstream or upstream of the gene that is responsible for production of their ligand. Changes in intracellular concentration of the ligand are sensed by the aptamer domain, leading to a conformational change of the expression platform and resulting in switching gene expression on or off (Nahvi et al., 2002; Winkler et al., 2002; Winkler et al., 2002). In eukaryotes, particularly in plants, algae, and fungi, intracellular thiamine pyrophosphate (TPP) levels are regulated by TPP-responsive riboswitches that function by alternative splicing of a TPP biosynthetic gene (Bocobza et al., 2007; Wachter et al., 2007; Wachter, 2014).

Inspired by natural riboswitches, researchers have created synthetic, ligand-responsive regulatory RNAs. To design synthetic riboswitches, mostly self-cleaving ribozymes are used as expression platforms that are linked to an aptamer domain via a communication sequence, yielding so-called aptazymes (ribozyme plus aptamer). Development of synthetic RNA aptamers via SELEX (Systematic Evolution of Ligands by Exponential enrichment) along with rational design of the communication sequences brought the possibility to generate aptazymes that sense a broad variety of molecular inputs, such as proteins, RNAs, metabolites and co-factors (Townshend et al., 2015; McKeague et al., 2016; Zhong et al., 2016). The option to induce conformational changes or destabilize mRNAs by a ligand-responsive aptazyme makes them versatile tools for genetic control in diverse biological systems. Both natural and synthetic riboswitches were used to regulate reporter and endogenous gene expression in a wide variety of organisms, including mammalian cells (Ausländer et al., 2010; Nomura et al., 2012; Beilstein et al., 2015), yeast (Win and Smolke, 2007; Wittmann and Suess, 2011; Klauser et al., 2015), plants (Bocobza et al., 2007; Wachter et al., 2007; Bocobza et al., 2013), algae (Ramundo et al., 2013), and cyanobacteria (Nakahira et al., 2013).

External control of gene expression is an important tool for biotechnology and the detailed analysis of gene functions in plants. In many cases, temporal or spatial control of transgene expression is needed to minimize disturbance of development of the plant or to avoid the presence of the gene product in non-target plant organs. To date, artificial inducible transcription factors (TFs), expressed either constitutively or from a tissue specific promoter, are the most broadly applied systems to achieve external induction of transgene expression in plants (Moore et al., 2006; Corrado and Karali, 2009). However, these systems have two major disadvantages: Firstly, they require the functionality of at least two transgenes, an inducer-sensitive transcription factor and the target gene with the binding site for the artificial transcription factor in its promoter region. Secondly, the expression level of artificial TFs may change substantially in response to endogenous (*e.g*. the cell cycle) or external factors. To repress an endogenous gene by an external trigger, inducible systems based on RNA interference (RNAi) were successfully employed (Guo et al., 2003; Ketelaar et al., 2004; Masclaux et al., 2004). In these systems, inducible expression of an RNA hairpin construct or an artificial micro-RNA (miRNA) activates the endogenous RNAi machinery resulting in the production of small interfering RNAs (siRNAs). The siRNAs subsequently mediate the transcriptional and posttranscriptional silencing of homologous genes (Borges and Martienssen, 2015). Another common approach to activate the RNAi machinery is virus-induced gene silencing (VIGS) that uses engineered plant viruses containing parts of the target gene(s) in their genome (Kumagai et al., 1995; Ruiz et al., 1998). All these systems rely on functionality and activity of the many components of the endogenous RNAi system. Additionally, plant viruses often have a limited host range and bear the risk of unwanted spreading from plant to plant (Senthil-Kumar and Mysore, 2011). Therefore, *cis*-acting systems like riboswitches, which allow combining the gene of interest and the regulatory element in a single transcript, promise to be significantly more robust and universal.

In the past decade, several studies were conducted to introduce riboswitch-mediated regulation of gene expression in plants (Bocobza et al., 2007; Wachter et al., 2007; Verhounig et al., 2010; Ogawa, 2011; Emadpour et al., 2015; Doron et al., 2016). The native TPP riboswitch from Arabidopsis (*Arabidopsis thaliana*) has been used successfully to regulate the expression of reporter genes (Bocobza et al., 2007; Wachter et al., 2007). However, the switching efficiency was lower than in transcription factor-based systems unless TPP-deficient mutants were used, which compromised the general suitability of this system for physiological analyses. Prokaryotic riboswitches and synthetic derivatives have been used to regulate expression of transgenes in the chloroplast genome of higher plants or microalgae (Verhounig et al., 2010; Emadpour et al., 2015; Doron et al., 2016). These systems allow very high expression levels and efficient ON or OFF-switching of transgene expression, but the subcellular localization of the proteins is limited to plastids. Regulation of nuclear-encoded genes in plants by synthetic riboswitches had not been reported so far.

Our approach to establish riboswitches as tools for regulation of nuclear gene expression in plants is based on a synthetic riboswitch that is responsive to an inducer not commonly found in the model plant Arabidopsis. Previous studies optimized theophylline-responsive riboswitches based on the self-cleaving hammerhead ribozyme (*HHR*) from *Shistosoma mansoni* for the regulation of gene expression in yeast and mammalian cells (Win and Smolke, 2007; Ausländer et al., 2010; Klauser et al., 2015; Rehm et al., 2015). The *HHR* in these switches can be completely inactivated by an A to G mutation in the catalytic core to demonstrate that the observed changes in gene expression depend on self-cleavage (Fig. 1A). To generate a ligand responsive aptazyme, stem-loop III of the *HHR* has been replaced by a synthetic theophylline aptamer linked via an optimized communication sequence (Yen et al., 2004; Ausländer et al., 2010). The aptamer used in our riboswitches was developed by *in vitro* evolution and has a high affinity and specificity for theophylline, a caffeine derivative (Jenison et al., 1994). For adoption of the active conformation of the catalytic core, *HHRs* depend on closure of stem III and tertiary interactions between loops L1 and L2 (Khvorova et al., 2003) (Fig. 1B). The proposed general base and general acid catalysis mechanism of the cleavage reaction involves interactions between one adenosine and two guanosine residues (Scott et al., 2013). When a ribozyme or aptazyme is inserted into the 3’-UTR of an mRNA, self-cleavage separates the poly-A-tail from the protein-coding sequence and thereby destabilizes the transcript (Fig. 1B). Incorporating the optimized theophylline-inducible aptazyme (*aTheoAz*) into the 3’-UTR of a luciferase reporter gene allowed reducing luciferase expression by more than 80% in HeLa cells (Ausländer et al., 2010).

**Figure 1.**
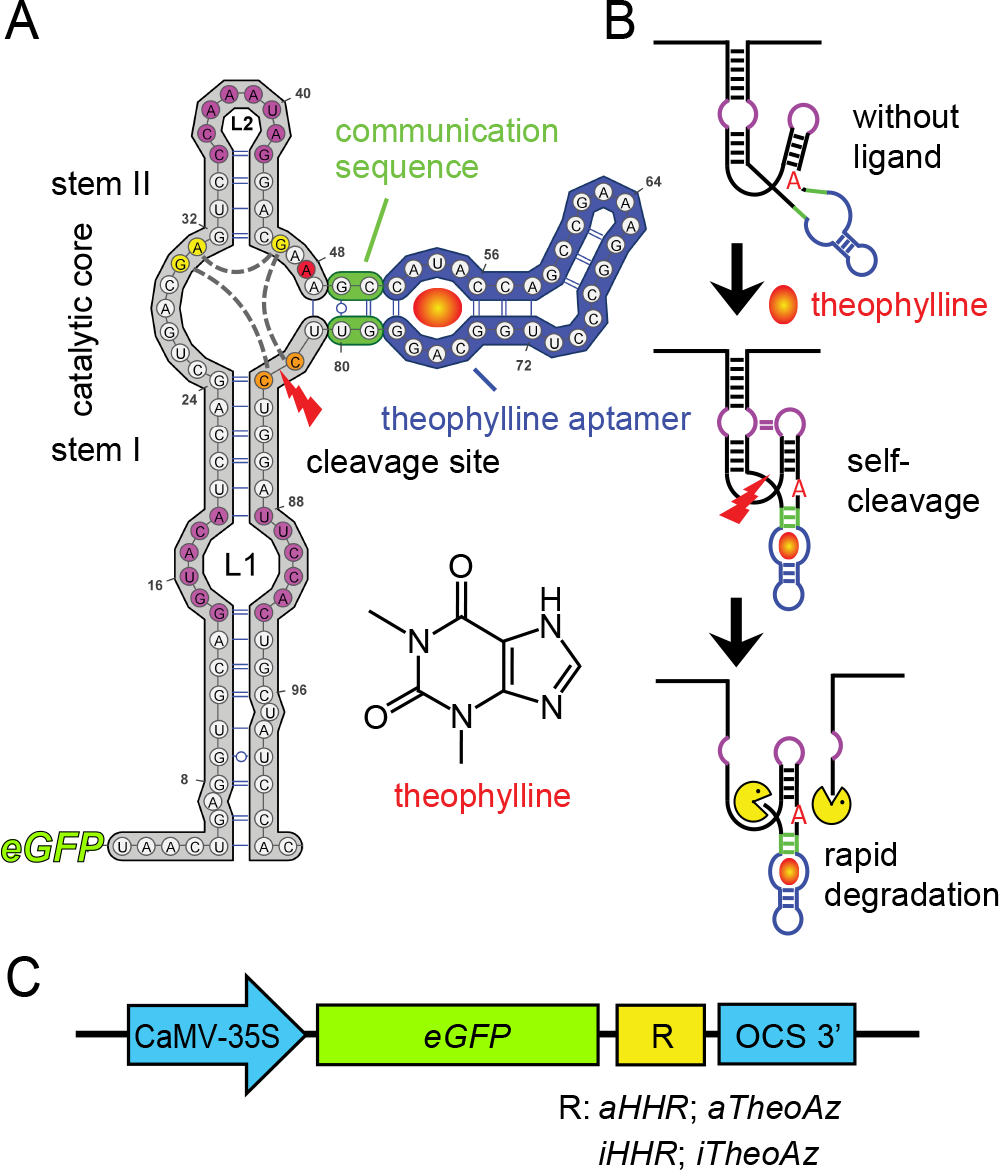
Structure and regulatory mechanism of the theophylline-inducible aptazyme. A, Sequence and secondary structure of the theophylline-inducible aptazyme (*aTheoAz*) built in VARNA (Darty et al., 2009). The hammerhead ribozyme (*HHR*) is plotted in grey, the communication sequence in green and the theophylline aptamer in blue. Tertiary interactions between loops L1 and L2 are indicated by magenta coloring. The A residue that is mutated to G in the inactive variants is marked in red. Nucleotides of the catalytic core that are involved in the cleavage reaction are marked in yellow, interactions between these nucleotides are indicated by grey dashed lines. The two cytosines between which the phosphodiester backbone is cleaved (cleavage site) are marked in orange. B, Proposed mechanism of de-stabilization of the *GFP* mRNA by theophylline-induced autocatalytic cleavage of the *aTheoAz*. C, Schematic illustration of the different constructs tested in transgenic Arabidopsis plants. *aHHR*: constitutively active *HHR*, *iHHR*: inactive *HHR*, *aTheoAz*: *aHHR* with theophylline aptamer, *iTheoAz*: *iHHR* with theophylline aptamer.

Here we show that the *aTheoAz* incorporated into the 3’-UTR of a *GFP* expression cassette constitutes an effective riboswitch enabling dose- and time-dependent down-regulation of GFP transcript and protein levels in Arabidopsis. We demonstrate that aptazyme-mediated downregulation of gene expression can be used for physiological studies by conditional complementation of mutants lacking the thylakoid-integral One Helix Protein 1 (OHP1), which is required for synthesis or maintenance of functional photosystems (Beck et al., 2017; Hey and Grimm, 2018; Myouga et al., 2018). The seedling-lethal phenotype of *ohp1-1* mutants was overcome by a switchable transgenic copy of the *OHP1* gene and treatment of seedlings with theophylline reduced the *OHP1* transcript level to near the detection limit. To our knowledge, this is the first report describing the use of a synthetic riboswitch to regulate nuclear-encoded genes *in planta*.

## RESULTS

### A library of ribozyme- and riboswitch-dependent expression cassettes

To analyze the functionality of the modified *S. mansoni* hammerhead ribozyme (*HHR*) and the theophylline-inducible aptazyme (*aTheoAz*) derived from it, we generated a series of different expression cassettes with the Green Fluorescent Protein (*GFP*) gene under control of the Cauliflower Mosaic Virus 35S promoter and the Octopine Synthase terminator (OCS-3’) from *Agrobacterium* TI-plasmids ((Ferbeyre et al., 1998), Fig. 1 and Supplemental Fig. S1). Between the coding sequence of *GFP* and the OCS-3’ we inserted either an active, modified *HHR* (*aHHR*) or an inactive variant (*iHHR*). Two other constructs contained either the *aTheoAz* or a theophylline-binding, but catalytically inactive aptazyme (*iTheoAz*; Fig. 1C). All constructs were stably inserted into Arabidopsis plants by *Agrobacterium*-mediated transformation and homozygous lines were selected for characterization.

### Suppression of gene expression by insertion of the hammerhead ribozyme into the 3’-UTR

Transgenic plants with the *GFP:aHHR* or *GFP:iHHR* constructs were screened for GFP expression by epifluorescence microscopy of leaves. As expected for free GFP, green fluorescence was observed in the cytosol and at slightly higher levels in the nuclei in lines with the *GFP:iHHR* construct. GFP fluorescence in lines with the *GFP:aHHR* construct was very low and is virtually invisible in images with gain and contrast settings optimized for the *GFP:iHHR* construct (Fig. 2A).

**Figure 2.**
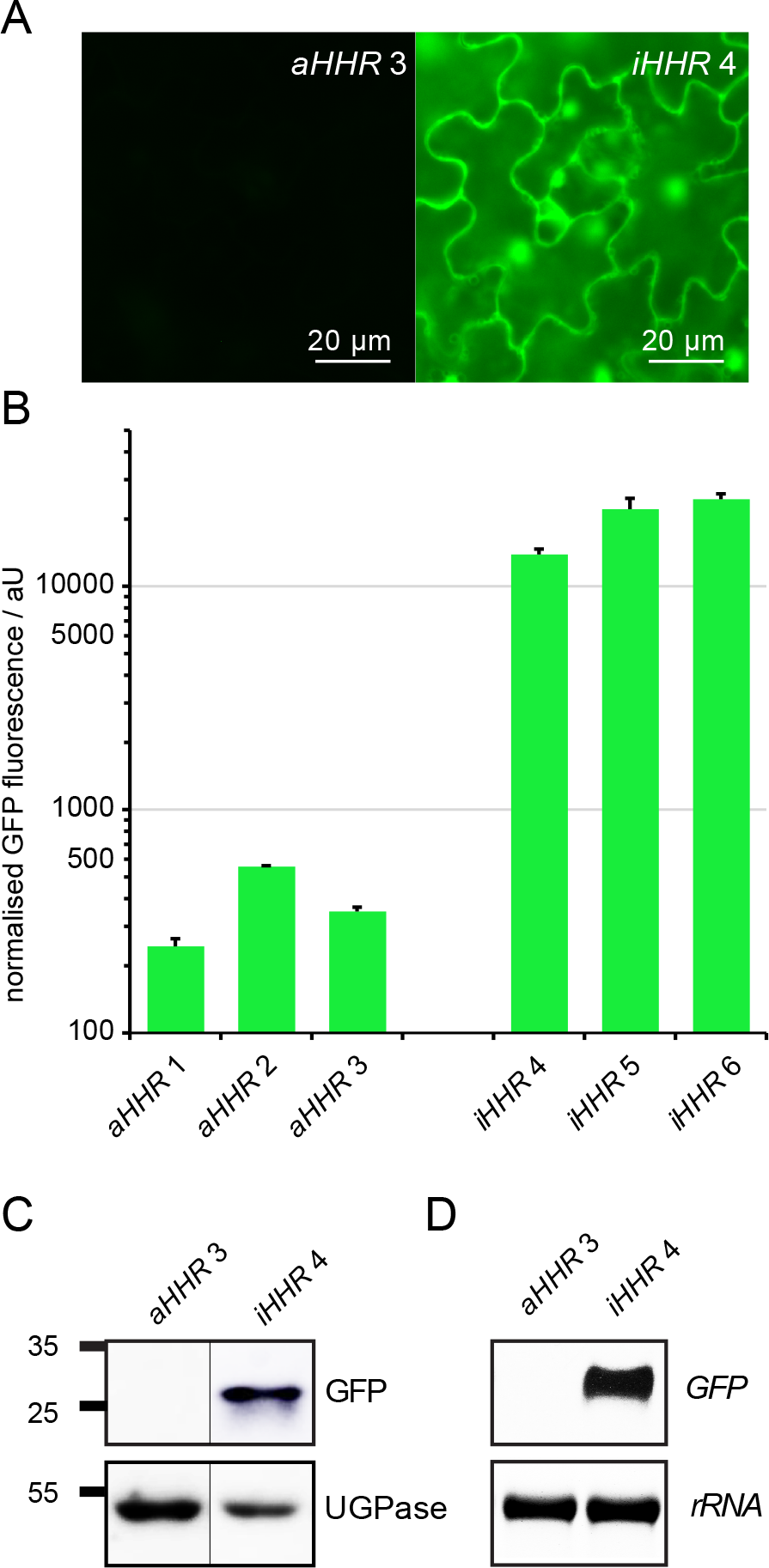
GFP expression levels in transgenic lines carrying the *GFP:aHHR* and *GFP:iHHR* constructs. A, Epifluorescence images of mature leaves from greenhouse-grown plants with the constitutively active ribozyme (*aHHR*) or the catalytically inactive (*iHHR*) ribozyme in the 3’-UTR of the *GFP* expression cassette. Exposure time and contrast settings are identical for both images. B, GFP fluorescence was quantified in soluble protein extracts of 2-week-old seedlings of independent transgenic lines grown under short-day conditions in sterile culture. Columns represent the average +SE of three independent biological replicates. C, Representative Western blot showing detection of GFP in soluble protein extracts of seedlings carrying the *GFP:iHHR* construct, but not in seedlings with the *GFP:aHHR* construct. UDP-glucose pyrophosphorylase (UGPase) was used as loading control, the positions of molecular weight markers are indicated at the left. D, Detection of *GFP* transcripts in 3-week-old seedlings by Northern blot. EtBr stained 25S-*rRNA* is shown as loading control.

To quantify the difference in GFP expression levels between lines carrying the active and inactive ribozymes, we analyzed GFP fluorescence in soluble protein extracts of 2-week-old seedlings. Lines with the *GFP:aHHR* construct showed very low levels of GFP fluorescence with a slight variation between independent transgenic lines, whereas lines with the *GFP:iHHR* construct mostly showed strong GFP expression with on average more than 50-fold higher fluorescence intensities (Fig. 2B). From the tested lines, we selected one representative homozygous line for each construct for the detection of GFP protein and mRNA by Western and Northern blot, respectively. While the *GFP:iHHR* construct induced high levels of GFP protein and transcript expression, neither GFP protein nor transcript were detected in transgenic plants with the *GFP:aHHR* construct when the exposure time was optimized for detection of GFP mRNA or protein in the *GFP:iHHR* plants (Fig. 2C,D).

### Downregulation of gene expression by the theophylline-inducible aptazyme inserted in the 3’-UTR

In order to determine the theophylline tolerance of Arabidopsis seedlings, we germinated wildtype seeds on agar plates with different concentrations of theophylline. After two weeks, the roots were shorter in all samples with addition of theophylline in comparison to the control samples without theophylline (Supplemental Fig. S2A). The root lengths in the control seedlings were 72.4 ± 3.0 mm and they were gradually reduced to 16.5 ± 1.2 mm in seedlings cultivated in the presence of 1.5 mM theophylline. Size and number of the leaves were also negatively correlated to the concentration of theophylline, but up to 1.5 mM all leaves appeared healthy without macroscopic chlorosis or necrosis. Chlorophyll fluorescence analysis revealed that at theophylline concentrations of 1 mM or more a progressive reduction of the quantum efficiency of photosystem II (Φ_PSII_) and inducible energy dissipation (NPQ) was occurring, whereas photodamage (lowering of Fv/Fm as measure of photosynthetic capacity) was negligible up to 1.5 mM theophylline (Supplemental Fig. S2B-D). Based on these experiments, we chose maximally 1.5 mM theophylline in the culture medium for detecting changes in GFP protein expression in transgenic plants carrying the *aTheoAz* or *iTheoAz* constructs.

In preliminary experiments, we observed that it takes up to two weeks until changes in GFP protein expression become apparent after the application of theophylline to axenically grown seedlings. Therefore, we cultivated transgenic plants carrying aptazymes on plates with 0 or 1 mM theophylline for 14 days for quantification of GFP expression by fluorescence measurements in soluble protein extracts. Three homozygous lines with the active aptazyme showed between 87% and 93% lower GFP expression upon growth on 1 mM theophylline compared to growth in the absence of theophylline (Supplemental Fig. S3A). In contrast, in all three homozygous lines with the inactive aptazyme a trend towards higher GFP expression after growth on 1 mM theophylline was observed, although the increase was only significant in one of the lines (Supplemental Fig. S3B).

From the tested lines, we selected one representative homozygous line for each construct and used these two lines for detailed characterization of theophylline-dependent control of GFP expression (Fig. 3). By fluorescence microscopy, a lower GFP signal intensity was detected in seedlings with the *GFP:aTheoAz* construct after two weeks cultivation in presence of 1.5 mM theophylline, whereas no decline in green fluorescence was observed in seedlings with the *GFP:iTheoAz* construct (Supplemental Fig. S4). Quantitative analysis of GFP fluorescence in soluble protein extracts showed that in the *GFP:aTheoAz* line 3, GFP expression was reduced by 88%, 92%, and 93% when the seedlings were cultivated for two weeks in the presence 0.5 mM, 1.0 mM, and 1.5 mM theophylline, respectively (Fig. 3A). Although the GFP fluorescence was not significantly different between 0.5 and 1.5 mM theophylline in the overall comparison, there was a strong negative correlation (Pearson coefficient −0.945) between theophylline concentration and GFP fluorescence. Similar results to the GFP fluorescence assay were obtained when GFP expression was quantified from Western blot signal intensities. The GFP signal intensity was reduced by 92%, 93%, and 95% when the seedlings were cultivated in the presence 0.5 mM, 1.0 mM, and 1.5 mM theophylline, respectively (Fig. 3B,C). In contrast, the GFP signal intensity in seedlings of the *GFP:iTheoAz* line 5 was increased 1.9-fold in the presence of 1 mM theophylline compared to the untreated control, while at 0.5 mM and 1.5 mM theophylline, the differences were not significant (Supplemental Fig. S3C).

**Figure 3.**
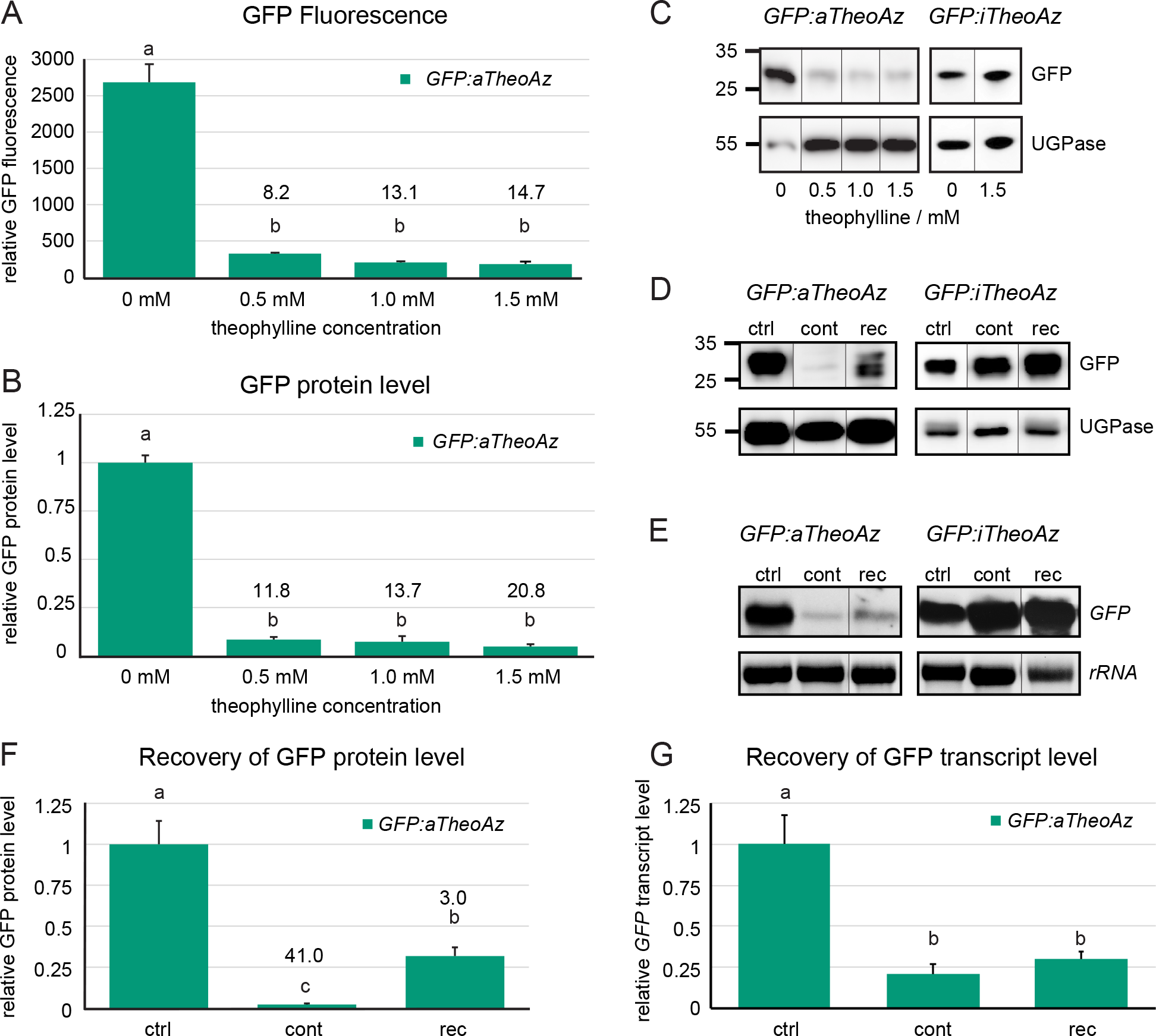
GFP expression levels in transgenic lines carrying the active or inactive theophylline aptazymes. A-C, Seedlings of *GFP:aTheoAz* line 3 or *GFP:iTheoAz* line 5 were grown on ½x MS-agar plates with 2 % Suc supplemented with 0 to 1.5 mM theophylline under long day conditions. After two weeks, soluble proteins were extracted and analyzed for GFP fluorescence and GFP protein level. A, Relative GFP fluorescence in soluble protein extracts. B, Relative GFP protein levels (normalized to UDP-glucose pyrophosphorylase and to control plants) in soluble protein extracts determined by Western blot. C, D, Representative Western blots of the quantitative analyses in B and F. Positions of molecular weight markers are indicated at the left. E, representative Northern blots of the quantitative analysis in G. F, G, Normalized GFP expression levels in extracts of seedlings cultivated for four weeks in the absence of theophylline (ctrl), for four weeks continuously on 1.5 mM theophylline (cont), or for 2 weeks on 1.5 mM theophylline followed by two weeks of recovery on plates without theophylline (rec) under long day conditions. F, GFP protein levels normalized to UGPase. G, *GFP* transcript levels normalized to 25S *rRNA*. In A, B, F and G, columns represent the average +SE of 3 biological replicates. Numbers above the bars represent the average response ratio (GFP fluorescence or expression level in control plants/GFP fluorescence or expression level in theophylline-treated plants). Different letters above the columns indicate significant differences (p<0.05 in one-way ANOVA with Tukey’s post-hoc Honestly Significant Difference test; Western blot signal intensities were square root-transformed for statistical analysis).

In order to analyze whether riboswitch-mediated down-regulation of gene expression is reversible, seedlings carrying the *GFP:aTheoAz* or *GFP:iTheoAz* constructs were grown for two weeks on medium with 1.5 mM theophylline and were then transferred to medium without theophylline. After two weeks of recovery, the level of GFP protein in the *GFP:aTheoAz* plants was 14 times higher compared to plants that were cultivated continuously in the presence of 1.5 mM theophylline (Fig. 3D,F). In seedlings exposed for 4 weeks to 1.5 mM theophylline, the level of GFP protein was reduced by 97% compared to control seedlings, whereas in the seedlings that were allowed to recover during two weeks, GFP expression was reduced only by 67%. When we analyzed GFP transcript levels in plants that were allowed to recover for two weeks from theophylline treatment, only a trend towards higher transcript levels but no significant recovery was observed (Fig. 3E,G). Under the same conditions, the presence or withdrawal of 1.5 mM theophylline did not alter GFP expression in the *GFP:iTheoAz* line (Supplemental Fig. S3D). To observe whether growth inhibition by theophylline is reversible, we also transferred WT plants to plates without theophylline after two weeks of growth on plates with 1.5 mM theophylline. While 4 weeks exposure to 1.5 mM theophylline under long-day conditions damaged the plants, the plants that were rescued after 2 weeks to plates without theophylline almost reached the same size as control plants (Supplemental Fig. S5).

To evaluate the short-term influence of theophylline on aptazyme-containing *GFP* transcripts, we performed Northern blots with total RNA extracted from mature leaves after application of theophylline or water via infiltration (Figure). When the infiltration was performed carefully, the leaves did not sustain mechanical injuries and did not develop lesions within 48 h after treatment with 2 mM theophylline (Supplemental Fig. S6). In leaves carrying the *iTheoAz* construct, high *GFP* transcript levels were detected that did not change 24 h after infiltration of leaves with 2 mM theophylline (Fig. 4A). To analyze the time- and dose-dependent response of the *GFP:aTheoAz* construct to theophylline application, we infiltrated leaves of transgenic line 3 with 0 to 2 mM theophylline and analyzed the level and integrity of *GFP* transcripts at different times after infiltration (Fig. 4A,B,C). Both full-length and truncated transcripts were detected in the leaves of plants with the *GFP:aTheoAz* construct even in water-infiltrated leaves (Fig. 4A). Three hours after infiltration with 2 mM theophylline, the reduction in the level of full-length *GFP* transcripts was the strongest and comprised 90.4% (corresponding to a 10.4-fold difference between the ON- and the OFF-state; Fig. 4B). Three hours after infiltration with 1.0 mM theophylline the reduction was 82.8% (corresponding to a 5.8-fold difference). Even in leaves infiltrated with 0.1 mM theophylline, the reduction in *GFP* transcript levels was significant 3 h after infiltration and comprised 52.4%. One and two days after infiltration with theophylline, the level of full-length *GFP* transcripts started to recover at most theophylline concentrations tested (Fig. 4C). The level of *GFP* transcripts 24 h and 48 h after infiltration with of 2 mM theophylline were reduced by 76.7% and 54.5% compared to the level of GFP transcripts water-infiltrated leaves. When the leaves were infiltrated with 0.4 mM theophylline or above, the reduction in *GFP* transcript levels was still significant at 24 h and 48 h after infiltration.

**Figure 4.**
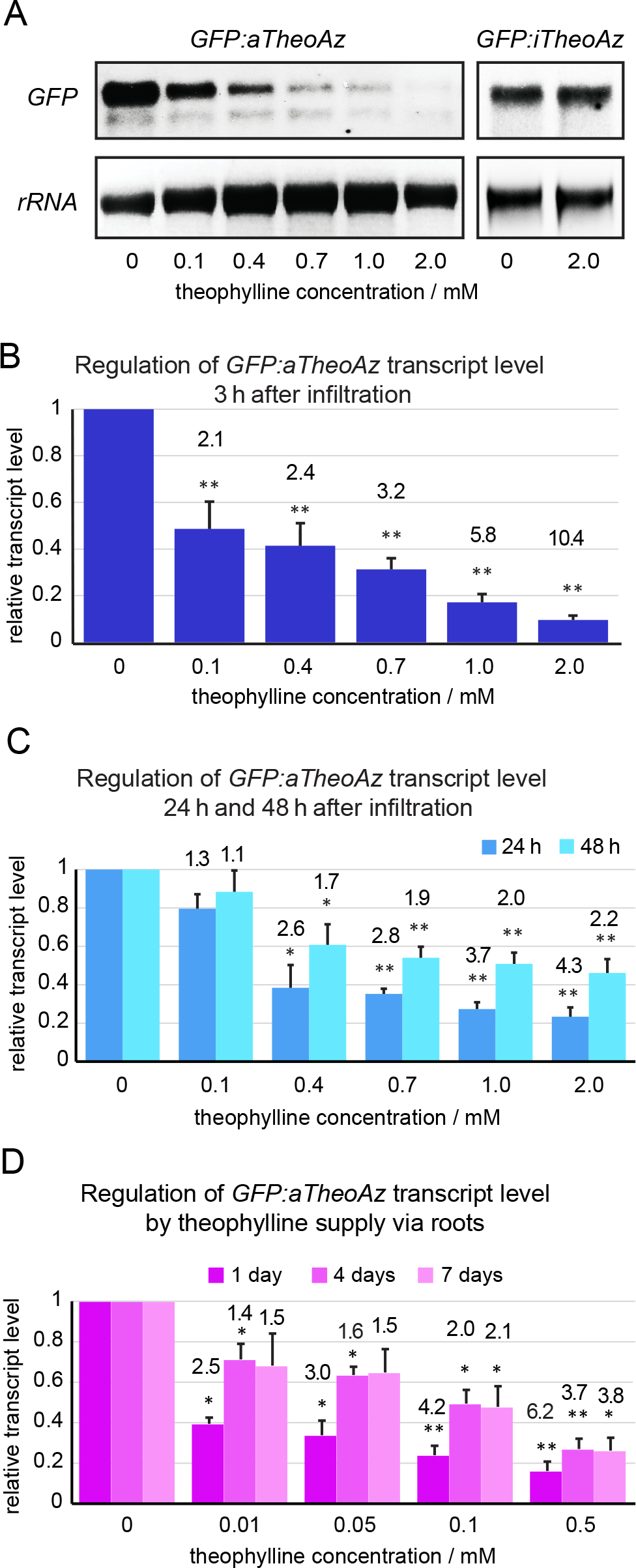
*GFP* transcript levels in transgenic lines carrying active and inactive theophylline aptazymes quantified by Northern blot analysis. A-C, Mature leaves of 4- to 6-week-old plants of *GFP:aTheoAz* line 3 were infiltrated with the indicated concentrations of theophylline and harvested at 3 h, 24 h or 48 h after infiltration. Leaves of plants from *GFP:iTheoAz* line 5 were harvested 24 h after infiltration with water or 2 mM theophylline. A, Representative Northern blots showing *GFP* transcripts of *GFP:aTheoAz* line 3 and *GFP:iTheoAz* line 5 at 3 h and 24 h, respectively, after application of 0 to 2 mM theophylline. B, C, Quantification of full-length *GFP* transcripts normalized to 25S *rRNA* and to the level of *GFP* transcripts in leaves infiltrated with water. Columns represent the average +SE from 5-7 Northern blots for each time series. Numbers above the columns represent the average response ratio (transcript level in water-treated leaves/transcript level in theophylline-infiltrated leaves). D, Three-week-old plants of *GFP:aTheoAz* line 3 were transferred to a hydroponic cultivation system (Supplemental Fig. 7). Two weeks after the transfer, the nutrient solution was supplemented with the indicated concentrations of theophylline. Full-length *GFP* transcript levels were quantified after 1 day, 4 days and 7 days of theophylline supply and normalized to the *25S rRNA* loading control and the samples without theophylline. Columns indicate the average +SE of 4 biological replicates, each containing leaves of two individual plants. In all panels, *, ** indicate significant differences to the water-infiltrated leaves (p < 0.05 and 0.01, respectively, in single sample T-tests of log-transformed response ratios).

Infiltration of leaves with theophylline solution offers the possibility to analyze theophylline- and water-infiltrated leaves from the same plant simultaneously but it requires a lot of manual work. For the observation of long-term effects of gene down-regulation, repeated infiltration would be required. Therefore, we analyzed the possibility to apply theophylline to mature plants via the roots. We chose a hydroponic cultivation system because soil microbes might alter the theophylline concentration in the growth medium. Three-week-old plants were transferred to a hydroponic cultivation system and after two weeks of further growth, the medium was supplemented with different concentrations of theophylline (Supplemental Fig. S7). At one, four and seven days after the addition of theophylline, the level of full-length *GFP* transcripts was quantified in leaf samples (Fig. 4D). At 0.01 mM theophylline, a significant drop of the *GFP* transcript level by 60.5% was evident after one day of theophylline application. With 0.1 mM theophylline or more, the *GFP* transcript levels were significantly lower than in the control plants at all analyzed time points. At all concentrations tested, the decline of the *GFP* transcript level was most pronounced one day after the onset of theophylline supply. After one day supply of 0.5 mM theophylline via the roots, the decrease of the *GFP* transcript level (84% less) was more pronounced compared to one day after infiltration of leaves with 0.4 mM theophylline (61.5% less; Fig. 4C,D). To avoid toxicity symptoms in our hydroponic cultivation system, which provides a virtually unlimited supply of theophylline with the constantly renewed nutrient solution, we did not test theophylline concentrations above 0.5 mM.

### *aTheoAz*-dependent downregulation of *OHP1* causes defects in photosynthesis

In order to be broadly applicable, an artificial riboswitch should be able to reduce gene expression of endogenous genes as well. Because an efficient method for site-specific and accurate insertion of foreign DNA into the genome of Arabidopsis is still missing, we chose to use conditional complementation of a T-DNA insertion mutant of *One Helix Protein 1* (*OHP1*) to demonstrate the utility of riboswitches for the regulation of endogenous genes. Failure to assemble or stabilize photosystems causes seedling lethality in *ohp1* mutants, which complicates the analysis of the precise gene function. A construct containing the *OHP1* cDNA with an *aTheoAz* in the 3’UTR under control of the CaMV 35S promoter had been used previously to complement the seedling-lethal phenotype of *ohp1-1* mutants (Beck et al., 2017). However, treatment of these plants with theophylline did not cause phenotypical changes, presumably because the residual transcript level still allowed the synthesis of sufficient amounts of OHP1 protein. Therefore, we generated a construct for switchable *OHP1* expression under the control of the endogenous promoter of *OHP1* and the theophylline aptazyme (*Pr_OHP1_:OHP1:aTheoAz*) and a corresponding construct containing an inactive aptazyme as control. When these constructs were introduced into heterozygous *ohp1-1* mutants, already in the T1-generation homozygous *ohp1-1* mutants were recovered that showed autotrophic growth and normal fertility. Most of the complemented mutants were visually indistinguishable from WT plants, and only such lines were used for further experiments.

We selected at least 3 homozygous lines (homozygous for both the *ohp1-1* mutation and the riboswitch construct) with either the active or the inactive aptazyme in the 3’-UTR of the *OHP1* expression cassette. Transfer of 2-week-old, axenically cultured seedlings to plates with 2 mM theophylline for four days caused a reduction of full-length *OHP1*-transcript levels by 78% to 90% in lines with the *aTheoAz* but not in lines with the *iTheoAz* construct (Fig. 5A). No phenotypic changes were visible in WT plants or complemented *ohp1-1* mutants under these conditions (Supplemental figure S8). When the mutants complemented with the *OHP1:aTheoAz* construct were germinated and cultivated in presence of 1.5 mM theophylline, the level of OHP1 protein was reduced by 76% in 4-week-old seedlings, whereas seedlings with the *OHP1:iTheoAz* construct had unchanged OHP1 levels (Fig. 5B). The decrease in OHP1 protein was accompanied by a chlorotic phenotype of the plants complemented with the *OHP1:aTheoAz* construct, resembling homozygous, uncomplemented *ohp1-1* mutants. In contrast, no significant changes in chlorophyll content were observed in lines with the *OHP1:iTheoAz* construct (Supplemental Fig. S9). Consistent with a reproduction of the *ohp1-1* mutant phenotype, photosynthetic capacity (Fv/Fm) was reduced in the lines with the *OHP1:aTheoAz* construct (Fig. 5C). Both chlorosis and reduction of Fv/Fm were most pronounced in the oldest leaves of the plants. In the absence of theophylline, photosynthetic capacity in the selected *OHP1:aTheoAz* line was 7.2% lower compared to Col-0 WT plants and *ohp1-1* mutants complemented with the *OHP1:iTheoAz* construct. Fv/Fm values were not changed by theophylline in Col-0 WT plants and in *ohp1-1* mutants complemented with the *OHP1:iTheoAz* construct. In contrast, Fv/Fm in seedlings of the *OHP1:aTheoAz* line was reduced by 23.2% compared to WT seedlings on plates with 1.5 mM theophylline. On plates without theophylline, all seedlings grew equally well, but in the presence of 1.5 mM theophylline, the *ohp1-1* mutants complemented with the *OHP1:aTheoAz* construct stayed smaller than WT seedlings and the *ohp1-1* seedlings with the *OHP1:iTheoAz* construct.

**Figure 5.**
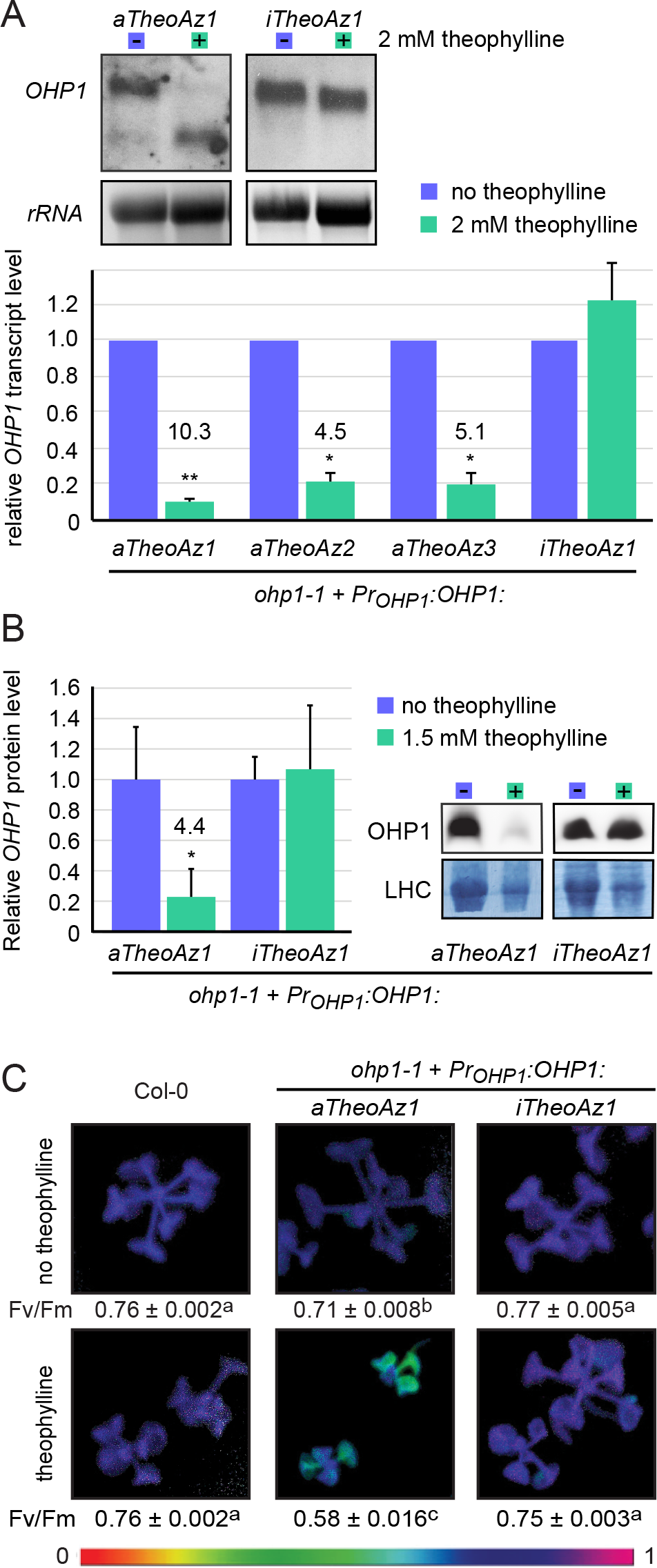
Aptazyme-mediated downregulation of *OHP1* expression. Homozygous *ohp1-1* seedlings carrying constructs with *OHP1* under control of its native promoter and either an active or an inactive aptazyme in the 3’-UTR were grown in axenic culture in short-day conditions. A, Two-week-old seedlings were transferred for four days to plates with or without 2 mM theophylline and *OHP1* transcripts in rosettes were detected by Northern blot. Panels: Representative blots showing *OHP1* transcript levels in mutants carrying the active riboswitch construct (*Pr_OHP1_:OHP1:aTheoAz*) or the inactive control construct (*Pr_OHP1_:OHP1:iTheoAz*) and EtBr stained 25S *rRNA* as loading control. Graph: Quantification of full-length *OHP1:aTheoAz* and *OHP1:iTheoAz* transcript levels normalized to the *rRNA* loading control and to control samples. B, OHP1 protein level in 4-week-old seedlings cultivated in the absence or presence of 1.5 mM theophylline. Panels: Representative blots with the Coomassie-stained membrane as loading control. Graph: Quantification of OHP1 signals relative to the LHC intensity. Columns in A and B represent the average +SE of three to four independent experiments. Numbers above the bars indicate the response ratios (expression levels in control samples/expression levels in treated samples). Significance was analyzed by single sample T-tests on log-transformed response ratios. *: p < 0.05 and **: p < 0.01. C, Photosynthetic capacity (Fv/Fm) of 4-week-old seedlings cultivated in the absence or presence of 1.5 mM theophylline. Chlorophyll fluorescence was measured before and during a single saturating flash and false-colored Fv/Fm images were generated with the ImagingWin software. Values below the pictures are the average ±SE of 11 to 17 seedlings. Different letters indicate significantly different values (p < 0.001 by two-way ANOVA with Tukey’s post-hoc Honestly Significant Difference test).

## DISCUSSION

In this study, we demonstrate the feasibility of using artificial aptazymes for the external regulation of nuclear-encoded genes in higher plants. As intended, the ligand-induced activation of the theophylline aptazyme (*aTheoAz*) derived from the *S. mansoni* hammerhead ribozyme allowed regulating the level of functional transcripts as well as the level of expressed protein for both, reporter and endogenous genes, with a high dynamic range. Independent of the target gene, activating the *aTheoAz* resulted in 84% to 95% downregulation of gene expression depending on theophylline concentration and application time.

When leaves of transgenic plants with the *aTheoAz* construct were infiltrated with theophylline solutions, full-length *GFP* transcripts rapidly disappeared in a dose-dependent manner. In leaves infiltrated with 0.1 mM theophylline, the *GFP* transcript level was reduced by 52% after 3 h, while after 24 h and 48 h, the effect was no longer significant. Twenty-four hours after infiltration with 0.4 mM or 0.7 mM theophylline, the *GFP* transcript levels were very similar compared to 3 h after infiltration. Even 48 h after infiltration, the *GFP* transcript level was still significantly decreased when theophylline concentrations of 0.4 mM or above were used. In leaves treated with 2 mM theophylline, the *GFP* transcript was reduced 3 h after infiltration to below 10% of the initial value, while it recovered to 23% after 24 h and to 46% after 48 h, indicating that theophylline is either diluted by transport to other parts of the plant or metabolized within the leaf. These results demonstrate that aptazyme-mediated transcript destabilization works rapidly and efficiently, and even after a single application of theophylline, there is a time window of at least 24 h in which the effect of lower transcript levels can be analyzed.

Because repeated infiltrations pose the risk of wounding stress, we added theophylline to the growth medium in sterile or hydroponic culture for prolonged, continuous exposure. Similar to the application via infiltration, adding theophylline to the hydroponic culture medium caused a transient decrease of the level of full-length *GFP* transcripts in plants carrying the *GFP:aTheoAz* construct. Even when only 0.01 mM theophylline were added to the nutrient solution, the level of *GFP* transcripts was reduced by 61% after one day. After transfer to medium with 0.5 mM theophylline, the reduction of the *GFP* transcript level by 84% after one day was very similar to leaves infiltrated with 1 mM theophylline (reduction by 83% after 3 h). This similar response to half the concentration of theophylline indicates that theophylline was efficiently taken up by the roots and transferred to the leaves, where it might have accumulated to higher concentrations than in the supplied nutrient solution. Despite the continuous supply of theophylline, the level of full-length *GFP* transcripts recovered partially after 4 and 7 days, supporting the hypothesis that theophylline can be metabolized in Arabidopsis leaves and the capacity to do so may be induced by theophylline exposure.

Growing Arabidopsis seedlings on sterile culture plates containing theophylline proved to be the most robust way to detect changes in GFP protein expression. Quantifying Western blot signals or measuring fluorescence in soluble protein extracts gave nearly identical results. In 2-week-old plants with the *GFP:aTheoAz* construct grown in the presence of 1.5 mM theophylline, GFP expression was reduced to below 5% of the level of plants grown in the absence of theophylline. In lines with the *GFP:iTheoAz* construct, GFP fluorescence never decreased in response to theophylline, on the contrary, a relatively small increase was observed occasionally. A previous report described symptoms of theophylline toxicity in tobacco at concentrations of 5 mM or above (Verhounig et al., 2010). We observed a dose-dependent inhibition of Arabidopsis seedling photosynthesis and growth on medium containing theophylline up to 1.5 mM, but symptoms of toxicity were observed only after 2 weeks on 2 mM or more theophylline. No negative effects were observed in leaves infiltrated once with 2 mM theophylline or in 3-week-old seedlings transferred to hydroponic nutrient solution containing up to 0.5 mM theophylline. Both, growth inhibition and downregulation of GFP expression were partially reversible. When 2-week-old seedlings with the *aTheoAz* construct grown on 1.5 mM theophylline were transferred to plates without theophylline for additional two weeks, the GFP protein level recovered to 33% of the level in the control seedlings, which was 14-times higher compared to seedlings that continued to grow on 1.5 mM theophylline (98% reduction compared to control seedlings). Under the same conditions, only a trend towards recovery of the *GFP* transcript level was observed, indicating that even small changes in the balance between translation and protein degradation can lead to the accumulation of higher levels of GFP protein. Either longer times would be needed to decrease intracellular theophylline to a concentration that allows significantly higher *GFP* transcript levels or the degradation products of the *GFP* transcripts had induced gene silencing. In seedlings with the *GFP:iTheoAz* construct, GFP expression stayed constant even after 4 weeks on 1.5 mM theophylline.

When the catalytically inactive hammerhead ribozyme (*iHHR*) was encoded in the 3’-UTR of a *GFP* overexpression cassette, only full-length transcripts were detected and GFP expression was high. Replacement of the *iHHR* with the active variant (*aHHR*) destabilized the mRNA and full-length transcripts as well as GFP protein were nearly undetectable. GFP fluorescence was reduced by 98% compared to the level detected in transgenic plants with the *iHHR* construct. Similar to the *iHHR*, insertion of the *iTheoAz* into the 3’UTR of the GFP expression cassette yielded high levels of GFP transcripts and protein that did not change consistently in response to theophylline. Plants carrying the *aTheoAz* in the 3’-UTR of the *GFP* mRNA showed on average 2- to 6-fold lower GFP expression levels compared to plants carrying the *GFP:iHHR* or *GFP:iTheoAz* constructs. Background activity in the absence of theophylline was also observed with the isolated *aTheoAz in vitro* and the high specificity of the theophylline aptamer makes it unlikely that endogenous metabolites interfere with the activation of the aptazyme (Jenison et al., 1994; Ausländer et al., 2010). The most likely explanation of the difference in *GFP* transcript levels between *GFP:aTheoAz* and *GFP:iTheoAz* lines is therefore the background activity of the aptazyme. This conclusion is supported by the detection of shorter *GFP* transcripts in the lines with the *GFP:aTheoAz* construct, which were not observed in lines with the *GFP:iTheoAz* construct. Despite the reduced basal GFP expression level, application of theophylline allowed downregulating GFP mRNA and protein levels by up to 95% in plants carrying the *GFP:aTheoAz* construct.

Currently, targeted genome editing in higher plants is limited to gene disruption whereas efficient and reliable methods for precise insertion of DNA constructs into the nuclear genome are still missing (Collonnier et al., 2017). Therefore, there is no straightforward possibility to regulate endogenous plant genes by introduction of artificial riboswitches. As a workaround for this challenge, we chose complementation of an insertion mutant to demonstrate the usefulness of riboswitches in the analysis or modulation of endogenous genes. Switchable copies of *OHP1*, either with the 35S promoter (Beck et al., 2017) or with the native *OHP1* promoter complemented the seedling-lethal, photosynthesis-deficient phenotype of homozygous *ohp1-1* mutants. Despite the much lower expression compared to the *GFP:aTheoAz* construct, theophylline treatment of plants with the *Pr_OHP1_:OHP1:aTheoAz* construct caused >90% reduction of transcript levels. When *ohp1-1* mutants complemented with the *OHP1:aTheoAz* construct under control of the native *OHP1* promoter were grown on plates with 1.5 mM theophylline, they stayed smaller than WT plants, had lower OHP1 protein levels, were slightly chlorotic, and showed a strong defect in photosynthetic capacity (Fv/Fm), which was not observed in complementation lines with the corresponding *OHP1:iTheoAz* construct. Fv/Fm values of *OHP1:aTheoAz* seedlings cultivated on 1.5 mM theophylline were 23% lower than in WT and *OHP1:iTheoAz* seedlings but still higher than in homozygous *ohp1-1* mutants (Beck et al., 2017), indicating that the residual translation of the *OHP1* mRNA before cleavage-induced degradation produced enough OHP1 protein to be partially functional. In another recent publication, no phenotypic alterations were observed after virus-induced gene silencing of *OHP1*, although *OHP1* mRNA levels were also reduced to approximately 10% of the level in control plants and OHP1 protein levels were strongly reduced (Hey and Grimm, 2018). Either the function of OHP1 was more important under our cultivation conditions or, more likely, riboswitch-mediated downregulation was more efficient during early developmental stages than virus-induced silencing. Additionally, the use of riboswitches allows the analysis of recovery after ligand withdrawal, whereas virus-induced silencing cannot readily be reversed.

Theophylline-inducible hammerhead aptazymes have previously been applied in *E. coli*, yeast and human cell lines. The observed changes of reporter gene expression were 10-fold upregulation in *E. coli* and downregulation by 75% to 83% in yeast and human cells in response to millimolar concentrations of theophylline (Wieland and Hartig, 2008; Ausländer et al., 2010; Klauser et al., 2015). The downregulation of GFP or OHP1 expression by 84% to 95% in our *aTheoAz-*carrying plants indicates that the effect of aptazyme-induced transcript destabilization caused a stronger downregulation of the protein level than in other model systems. Hence, we conclude that ligand-inducible aptazymes can be used effectively as artificial genetic switches in plants. Alternative systems based on the inducible expression of RNAi constructs or artificial microRNAs achieved reductions of transcript levels by 60% to 85% (O’Maoileidigh et al., 2015; Liu and Yoder, 2016; Thomson et al., 2017). However, in some cases, no efficient silencing was observed and the targeted genes tended to remain silenced after inducer withdrawal (O’Maoileidigh et al., 2015; Liu and Yoder, 2016). Because RNAi-based systems for gene silencing involve the generation and amplification of siRNAs by the targeted cells, they are difficult to tune. In contrast, the cleavage speed of artificial aptazymes and thereby the lifetime of the mRNA in which they reside is directly controlled by the ligand concentration. Therefore, artificial riboswitches constitute a promising addition to the molecular toolbox for basic research and biotechnology in plants. They may even be more robust than transcription factor-based systems because they act primarily in *cis* and do not rely on functional interactions between multiple transgenic elements.

Alternative ligands (*e.g.* TPP, guanine) for artificial riboswitches are available and several further aptamers are currently under development (Wieland et al., 2009; Wittmann and Suess, 2011; Nomura et al., 2012; Beilstein et al., 2015; Lotz and Suess, 2018). TPP application was previously used to regulate the expression of *GFP*, *YFP* or firefly *luciferase* fused to the 3’-UTR of the *THIC* gene containing the natural TPP riboswitch (Bocobza et al., 2007; Wachter et al., 2007). Bocobza *et al.* (2007) observed efficient switching only in a thiamine-deficient mutant, whereas Wachter *et al.* (2007) observed a TPP-dependent reduction of GFP fluorescence by approximately 50% in wildtype background. In our artificial switch with a ligand that is not produced in Arabidopsis, the dynamic range was much greater. With the recent development of ribozyme-based ON-switches for eukaryotic gene expression, the range of possible applications will be significantly increased (Beilstein et al., 2015; Wurmthaler et al., 2019). Especially for endogenous pest control in crop plants, temporally or spatially limited expression of effector molecules is very important to impede the formation of resistances in the pathogens or herbivores (Bates et al., 2005). Additionally, the insertion of switches into more than one transgene would allow differential regulation of two or more genes by a single treatment. The use of artificial ribozymes in crop biotechnology or for the controlled production of pharmaceuticals in engineered plants or cell cultures will require the development of novel aptazymes with ligands that do not affect plant metabolism and that are non-hazardous for humans and for the environment. As more and more structural data on riboswitches and ligand-binding RNA aptamers becomes available, also the targeted combination of aptamers and alternative ribozymes or other functional domains to form novel aptazymes will be facilitated (Lotz and Suess, 2018).

## CONCLUSIONS

The theophylline-dependent aptazyme described in this study can be used as a tunable genetic off-switch in the analysis of nuclear transgenes or complementation constructs in Arabidopsis and most likely in any plant species that does not produce theophylline. By the use of the inactive variant of the aptazyme as control, the specific effect of mRNA and protein depletion can be reliably separated from potential side effects of theophylline application. When short-lived proteins or RNAs are investigated, infiltration of mature leaves with theophylline solutions up to 2 mM provides a rapid system to study the specific effects of RNA destabilization and subsequent downregulation of protein levels. For the depletion of more stable proteins, we propose continuous application of up to 1.5 mM theophylline via the growth medium in hydroponic culture or in axenic culture. With these three methods of theophylline application, the effect of target protein depletion can be analyzed at virtually any developmental stage of the plant.

## MATERIAL AND METHODS

### Plant material and growth conditions

*Arabidopsis thaliana* (L) Heynh. ecotype Columbia-0 was obtained from the NASC (Stock-Nr. N60000) and *ohp1-1* T-DNA insertion mutants (GABI_362D02) were obtained from the GABI-KAT project (Kleinboelting et al., 2012; Beck et al., 2017). Plants were grown on soil in greenhouse chambers with either short-day (9 h/21 °C light, 15 h/17 °C darkness) or long-day (16 h/21 °C light, 8 h/17 °C darkness) conditions at 120±30 μmol photons m^−2^ s^−1^ at 60% relative humidity unless otherwise stated. For segregation analysis, protein extraction and for chlorophyll fluorescence measurement, surface-sterilized seeds were plated on Petri dishes with ½x Murashige and Skoog (MS) medium supplemented with 2% (w/v) sucrose and solidified with 0.8% (w/v) agar. The plates were kept under short-day or long-day conditions at 21 °C and 100±15 μmol photons m^−2^ s^−1^. For hydroponic culture, 3-week-old seedlings grown on soil were transferred to a hydroponic cultivation system in a climate chamber with short-day conditions. Eight plants each shared a pot with 2 l nutrient solution (1 mM Ca(NO_3_)_2_, 500 μM MgSO_4_, 500 μM K_2_HPO_4_, 100 μM KCl, 20 μM Fe-EDDHA, 10 μM H_3_BO_3_, 0.5 μM NiSO_4_, 0.2 μM Na_2_MoO_4_, 0.1 μM MnSO_4_, 0.1 μM CuSO_4_, 0.1 μM ZnSO_4_, 1 mM MES pH 6.0; adapted from (Küpper et al., 2007). The nutrient solution was aerated with an aquarium pump and renewed at a rate of 1 l/day.

### Theophylline application

Theophylline was dissolved in water at a concentration of 20 mM and sterilized by filtration. This stock was used to supplement the cultivation medium or to generate working solutions for leaf infiltration. Disposable 1 ml syringes gently pressed to the lower epidermis were used to infiltrate individual leaves of greenhouse-grown plants.

### Plasmid constructs and plant transformation

Ribozymes or theophylline aptazymes were amplified by PCR and inserted into the 3′-UTR of *eGFP* derived from the binary vector pEZT-NL using EcoRI and KpnI sites (Erhardt). The promoterless expression cassettes including the OCS terminator were subcloned in pENTR-D-Topo and transferred into pEG100 (Earley et al., 2006) by LR recombination (ThermoFisher). The CDS of *OHP1* and the native *OHP1* promoter (593 bp upstream of the start codon) were inserted by Gibson assembly (TaKaRa, NEB) into the above-mentioned constructs by replacing *GFP* and the CaMV 35S promoter consecutively. *Agrobacterium tumefaciens* strain GV3101 was used to introduce the constructs into wild type plants or heterozygous *ohp1-1* mutants by floral dip (Clough and Bent, 1998). Transgenic plants were selected by spraying with 50 μg ml^−1^ BASTA and the presence of the T-DNA was verified by PCR. Segregation analysis in the T2 and T3 generations was used to identify homozygous plants of single-insertion lines.

### RNA isolation and Northern Blot

Total RNA was isolated from deep-frozen plant tissue using commercial guanidine thiocyanate/phenol reagent (Chomczynski, 1993). Northern blots were carried out using the DIG labeling and detection system (Roche). Per lane, 10 μg (for *GFP*) or 15 μg (for *OHP1*) of total RNA were loaded. RNA separation, transfer to a nylon membrane, hybridization and detection were performed as described by (Woitsch and Romer, 2003). DIG-labeled probes comprising the entire CDS of *OHP1* or *GFP* were generated by PCR, column-purified and diluted in high-SDS hybridization buffer. Signal intensities on scanned X-ray films or digital images were determined with Image J. Relative numbers for gene expression were obtained by normalizing the specific signal intensity with the intensity of the EtBR stained 25S *rRNA*.

### Protein extraction and analysis

Fresh leaf tissue or entire seedlings were mixed with 10 μl mg^−1^ protein extraction buffer (50 mM Tris-HCl pH 8, 50 mM KCl, 50 mM MgCl_2_) in 2 ml safelock tubes and homogenized with a single steel bead for 3 min at 20 Hz in a TissueLyser (Qiagen). After centrifugation for 3 min at 26000 g at 4 °C, 200 μl of supernatant was transferred into a 96-well plate with black walls and thin, transparent, flat bottoms in a 5-step series of 1:2 dilutions. GFP fluorescence was measured from the top in a plate reader (Tecan) with excitation at 485 nm and emission detection at 535 nm. OD_280_ was recorded as a measure of total protein concentration. The region of linear correlation between dilution and fluorescence or extinction was determined visually and the slope of a linear regression was used as measure for fluorescence intensity or protein concentration. Green fluorescence was normalized to total protein content and the autofluorescence of wildtype protein extract was subtracted to obtain normalized GFP expression levels. For Western blotting, the soluble protein extract was mixed with lithium dodecyl sulphate loading buffer and 20 μg of protein per lane were separated on discontinuous 12.5% polyacrylamide gels. For OHP1 detection, insoluble proteins were separated on 15% Tris-tricine gels. Transfer to PVDF membrane and detection with polyclonal rabbit anti-GFP (Chromotek PABG1), anti-UDP-glucose pyrophosphorylase (Agrisera AS05 086), or anti-OHP1 (Hey and Grimm, 2018) antibodies and HRP-coupled goat anti-rabbit secondary antibody followed standard procedures.

### Pulse amplitude modulated (PAM) fluorimetry and chlorophyll quantification

Chlorophyll fluorescence was monitored using an Imaging PAM Fluorimeter (Walz) equipped with a standard measuring head using the Imaging Win software provided by the supplier. Program settings were: measuring light intensity 1, measuring light frequency 1, damping 2, gain 9, saturating pulse intensity 10, actininc light intensity 5, yield filter 3 and Fm-factor 1.024. Before the measurements, whole plants were dark-adapted for 5 min. Photosynthetic capacity (Fv/Fm) of homogenously illuminated areas of selected leaves was calculated from images recorded before and after the first saturating flash. Then the plants were illuminated with 80 μmol sec^−1^ m^−2^ blue light and further saturating flashes were used to monitor the induction of energy dissipation in the antenna (NPQ) and the operating efficiency of photosystem II (Φ_PSII_) which typically reached a steady-state level after 5 min. Chlorophyll was extracted by homogenizing fresh seedlings in 9 μl*mg^−1^ of 88% acetone. Absorption was determined in appropriate dilutions with 80% acetone at 470 nm, 646.8 nm, and 663.2 nm and chlorophyll content was calculated according to (Lichtenthaler, 1987).

## ACKNOWLEDGEMENTS

We thank Roswitha Miller and Silvia Kuhn for technical support and the gardeners of the botanical garden of the University of Konstanz for excellent plant care. Marc Stift is acknowledged for help with the statistical analysis. We thank Bernhard Grimm for providing the OHP1 antibody and Erika Isono for constructive comments on the manuscript.

## REFERENCES

Ausländer S, Ketzer P, Hartig JS (2010) A ligand-dependent hammerhead ribozyme switch for controlling mammalian gene expression. Mol Biosyst 6: 807–814

Bates SL, Zhao JZ, Roush RT, Shelton AM (2005) Insect resistance management in GM crops: past, present and future. Nat Biotechnol 23: 57–62

Beck J, Lohscheider JN, Albert S, Andersson U, Mendgen KW, Rojas-Stütz MC, Adamska I, Funck D (2017) Small One-Helix Proteins Are Essential for Photosynthesis in Arabidopsis. Front Plant Sci 8: 7

Beilstein K, Wittmann A, Grez M, Suess B (2015) Conditional control of mammalian gene expression by tetracycline-dependent hammerhead ribozymes. ACS Synth Biol 4: 526–534

Bocobza S, Adato A, Mandel T, Shapira M, Nudler E, Aharoni A (2007) Riboswitch-dependent gene regulation and its evolution in the plant kingdom. Genes Dev 21: 2874–2879

Bocobza SE, Malitsky S, Araujo WL, Nunes-Nesi A, Meir S, Shapira M, Fernie AR, Aharoni A (2013) Orchestration of thiamin biosynthesis and central metabolism by combined action of the thiamin pyrophosphate riboswitch and the circadian clock in Arabidopsis. Plant Cell 25: 288–307

Borges F, Martienssen RA (2015) The expanding world of small RNAs in plants. Nature reviews. Molecular cell biology 16: 727–741

Chomczynski P (1993) A reagent for the single-step simultaneous isolation of RNA, DNA and proteins from cell and tissue samples. BioTechniques 15: 532-534, 536-537

Clough SJ, Bent AF (1998) Floral dip: a simplified method for Agrobacterium-mediated transformation of *Arabidopsis thaliana*. Plant J 16: 735–743

Collonnier C, Guyon-Debast A, Maclot F, Mara K, Charlot F, Nogue F (2017) Towards mastering CRISPR-induced gene knock-in in plants: Survey of key features and focus on the model Physcomitrella patens. Methods 121-122: 103–117

Corrado G, Karali M (2009) Inducible gene expression systems and plant biotechnology. Biotechnol Adv 27: 733–743

Darty K, Denise A, Ponty Y (2009) VARNA: Interactive drawing and editing of the RNA secondary structure. Bioinformatics (Oxford, England) 25: 1974–1975

Doron L, Segal N, Shapira M (2016) Transgene Expression in Microalgae - From Tools to Applications. Front Plant Sci 7: 505

Earley KW, Haag JR, Pontes O, Opper K, Juehne T, Song K, Pikaard CS (2006) Gateway-compatible vectors for plant functional genomics and proteomics. Plant J 45: 616–629

Emadpour M, Karcher D, Bock R (2015) Boosting riboswitch efficiency by RNA amplification. Nucleic Acids Res 43: e66

Erhardt D Carnegie Cell Imaging Project (http://deepgreen.stanford.edu)

Ferbeyre G, Smith JM, Cedergren R (1998) Schistosome satellite DNA encodes active hammerhead ribozymes. Mol Cell Biol 18: 3880–3888

Guo HS, Fei JF, Xie Q, Chua NH (2003) A chemical-regulated inducible RNAi system in plants. Plant J 34: 383–392

Hey D, Grimm B (2018) ONE-HELIX PROTEIN2 (OHP2) Is Required for the Stability of OHP1 and Assembly Factor HCF244 and Is Functionally Linked to PSII Biogenesis. Plant Physiol 177: 1453–1472

Jenison RD, Gill SC, Pardi A, Polisky B (1994) High-resolution molecular discrimination by RNA. Science 263: 1425–1429

Ketelaar T, Allwood EG, Anthony R, Voigt B, Menzel D, Hussey PJ (2004) The actin-interacting protein AIP1 is essential for actin organization and plant development. Curr Biol 14: 145–149

Khvorova A, Lescoute A, Westhof E, Jayasena SD (2003) Sequence elements outside the hammerhead ribozyme catalytic core enable intracellular activity. Nat Struct Biol 10: 708–712

Klauser B, Atanasov J, Siewert LK, Hartig JS (2015) Ribozyme-based aminoglycoside switches of gene expression engineered by genetic selection in S. cerevisiae. ACS Synth Biol 4: 516–525

Klauser B, Rehm C, Summerer D, Hartig JS (2015) Engineering of ribozyme-based aminoglycoside switches of gene expression by in vivo genetic selection in Saccharomyces cerevisiae. Methods Enzymol 550: 301–320

Kleinboelting N, Huep G, Kloetgen A, Viehoever P, Weisshaar B (2012) GABI-Kat SimpleSearch: new features of the Arabidopsis thaliana T-DNA mutant database. Nucleic Acids Res 40: D1211–1215

Kumagai MH, Donson J, della-Cioppa G, Harvey D, Hanley K, Grill LK (1995) Cytoplasmic inhibition of carotenoid biosynthesis with virus-derived RNA. Proc Natl Acad Sci U S A 92: 1679–1683

Küpper H, Parameswaran A, Leitenmaier B, Trtilek M, Setlik I (2007) Cadmium-induced inhibition of photosynthesis and long-term acclimation to cadmium stress in the hyperaccumulator Thlaspi caerulescens. New Phytol 175: 655–674

Lichtenthaler HK (1987) Chlorophylls and carotenoids: Pigments of photosynthetic biomembranes. In Methods Enzymol, Vol Volume 148. Academic Press, p 350

Liu S, Yoder JI (2016) Chemical induction of hairpin RNAi molecules to silence vital genes in plant roots. Sci Rep 6: 37711

Lotz TS, Suess B (2018) Small-Molecule-Binding Riboswitches. Microbiol Spectr 6

Masclaux F, Charpenteau M, Takahashi T, Pont-Lezica R, Galaud JP (2004) Gene silencing using a heat-inducible RNAi system in Arabidopsis. Biochem Biophys Res Commun 321: 364–369

McKeague M, Wong RS, Smolke CD (2016) Opportunities in the design and application of RNA for gene expression control. Nucleic Acids Res 44: 2987–2999

Mellin JR, Cossart P (2015) Unexpected versatility in bacterial riboswitches. Trends Genet 31: 150–156

Moore I, Samalova M, Kurup S (2006) Transactivated and chemically inducible gene expression in plants. Plant J 45: 651–683

Myouga F, Takahashi K, Tanaka R, Nagata N, Kiss AZ, Funk C, Nomura Y, Nakagami H, Jansson S, Shinozaki K (2018) Stable Accumulation of Photosystem II Requires ONE-HELIX PROTEIN1 (OHP1) of the Light Harvesting-Like Family. Plant Physiol 176: 2277–2291

Nahvi A, Sudarsan N, Ebert MS, Zou X, Brown KL, Breaker RR (2002) Genetic control by a metabolite binding mRNA. Chem Biol 9: 1043

Nakahira Y, Ogawa A, Asano H, Oyama T, Tozawa Y (2013) Theophylline-dependent riboswitch as a novel genetic tool for strict regulation of protein expression in Cyanobacterium Synechococcus elongatus PCC 7942. Plant Cell Physiol 54: 1724–1735

Nomura Y, Kumar D, Yokobayashi Y (2012) Synthetic mammalian riboswitches based on guanine aptazyme. Chem Commun (Camb) 48: 7215–7217

O’Maoileidigh DS, Thomson B, Raganelli A, Wuest SE, Ryan PT, Kwasniewska K, Carles CC, Graciet E, Wellmer F (2015) Gene network analysis of *Arabidopsis thaliana* flower development through dynamic gene perturbations. Plant J 83: 344–358

Ogawa A (2011) Rational design of artificial riboswitches based on ligand-dependent modulation of internal ribosome entry in wheat germ extract and their applications as label-free biosensors. RNA 17: 478–488

Ramundo S, Rahire M, Schaad O, Rochaix JD (2013) Repression of essential chloroplast genes reveals new signaling pathways and regulatory feedback loops in chlamydomonas. Plant Cell 25: 167–186

Rehm C, Klauser B, Hartig JS (2015) Engineering aptazyme switches for conditional gene expression in mammalian cells utilizing an in vivo screening approach. Methods Mol Biol 1316: 127–140

Ruiz MT, Voinnet O, Baulcombe DC (1998) Initiation and maintenance of virus-induced gene silencing. Plant Cell 10: 937–946

Scott WG, Horan LH, Martick M (2013) The hammerhead ribozyme: structure, catalysis, and gene regulation. Prog Mol Biol Transl Sci 120: 1–23

Senthil-Kumar M, Mysore KS (2011) New dimensions for VIGS in plant functional genomics. Trends Plant Sci 16: 656–665

Sherwood AV, Henkin TM (2016) Riboswitch-Mediated Gene Regulation: Novel RNA Architectures Dictate Gene Expression Responses. Annu Rev Microbiol 70: 361–374

Thomson B, Graciet E, Wellmer F (2017) Inducible Promoter Systems for Gene Perturbation Experiments in Arabidopsis. Methods Mol Biol 1629: 15–25

Townshend B, Kennedy AB, Xiang JS, Smolke CD (2015) High-throughput cellular RNA device engineering. Nat Methods 12: 989–994

Verhounig A, Karcher D, Bock R (2010) Inducible gene expression from the plastid genome by a synthetic riboswitch. Proc Natl Acad Sci U S A 107: 6204–6209

Wachter A (2014) Gene regulation by structured mRNA elements. Trends Genet 30: 172–181

Wachter A, Tunc-Ozdemir M, Grove BC, Green PJ, Shintani DK, Breaker RR (2007) Riboswitch control of gene expression in plants by splicing and alternative 3′ end processing of mRNAs. Plant Cell 19: 3437–3450

Wieland M, Benz A, Klauser B, Hartig JS (2009) Artificial ribozyme switches containing natural riboswitch aptamer domains. Angew Chem Int Ed Engl 48: 2715–2718

Wieland M, Hartig JS (2008) Improved aptazyme design and in vivo screening enable riboswitching in bacteria. Angew Chem Int Ed Engl 47: 2604–2607

Win MN, Smolke CD (2007) A modular and extensible RNA-based gene-regulatory platform for engineering cellular function. Proc Natl Acad Sci U S A 104: 14283–14288

Winkler W, Nahvi A, Breaker RR (2002) Thiamine derivatives bind messenger RNAs directly to regulate bacterial gene expression. Nature 419: 952–956

Winkler WC, Cohen-Chalamish S, Breaker RR (2002) An mRNA structure that controls gene expression by binding FMN. Proc Natl Acad Sci U S A 99: 15908–15913

Wittmann A, Suess B (2011) Selection of tetracycline inducible self-cleaving ribozymes as synthetic devices for gene regulation in yeast. Mol Biosyst 7: 2419–2427

Woitsch S, Romer S (2003) Expression of xanthophyll biosynthetic genes during light-dependent chloroplast differentiation. Plant Physiol 132: 1508–1517

Wurmthaler LA, Sack M, Gense K, Hartig JS, Gamerdinger M (2019) A tetracycline-dependent ribozyme switch allows conditional induction of gene expression in Caenorhabditis elegans. Nat Commun 10: 491

Yen L, Svendsen J, Lee JS, Gray JT, Magnier M, Baba T, D’Amato RJ, Mulligan RC (2004) Exogenous control of mammalian gene expression through modulation of RNA self-cleavage. Nature 431: 471–476

Zhong G, Wang H, Bailey CC, Gao G, Farzan M (2016) Rational design of aptazyme riboswitches for efficient control of gene expression in ammalian cells. Elife 5

